# Ambulatory physiological measures obtained under naturalistic urban mobility conditions have acceptable reliability

**DOI:** 10.1101/2024.04.26.590892

**Authors:** Dilber Korkmaz, Kilian Knauth, Angela Brands, Marie Schmeck, Pia Büning, Jan Peters

## Abstract

Ambulatory assessment methods in psychology and clinical neuroscience are powerful research tools for collecting data outside of the laboratory. These methods encompass physiological, behavioral, and self-report measures obtained while individuals navigate in real-world environments, thereby increasing the ecological validity of experimental approaches. Despite the recent increase in applications of ambulatory physiology, data on the reliability of these measures is still limited. To address this issue, twenty-six healthy participants (*N* = 15 female, 18-34 years) completed an urban walking route (2.1 km, 30 min walking duration, temperature *M* = 19.8° degree Celsius, *Range* = 12°-37° degrees Celsius) on two separate testing days, while GPS-location and ambulatory physiological measures (cardiovascular and electrodermal activity) were continuously recorded. Bootstrapped test-retest reliabilities of single measures and aggregate scores derived via principal component analysis (PCA) were computed. The first principal component (PC#1) accounted for 39% to 45% of variance across measures. PC#1 scores demonstrated an acceptable test-retest reliability (*r* = .60) across testing days, exceeding the reliabilities of most individual measures (heart rate: *r* = .53, heart rate variability: *r* = .50, skin conductance level: *r* = .53, no. of skin conductance responses: *r* = .28, skin conductance response amplitude: *r* = .60). Results confirm that ambulatory physiological measures recorded during naturalistic navigation in urban environments exhibit acceptable test-retest reliability, in particular when compound scores across physiological measures are analyzed, a prerequisite for applications in (clinical) psychology and digital health.

**Author summary:** Psychophysiological assessments have been predominantly limited to controlled laboratory settings, leaving the reliability of field measurements unclear. In this study, we conducted a proof-of-concept investigation in *N*=26 healthy participants navigating the same urban route on two separate days. Cardiovascular and electrodermal activity were continuously recorded and combined with GPS-based location tracking. Psychophysiological measurements obtained under naturalistic urban mobility conditions showed acceptable test-retest reliability, in particular when multiple measures where combined into a compound score via principal component analysis. Shedding light on the reliability of ambulatory assessments in urban environments emphasizes the potential for psychophysiological measurements to contribute valuable insights beyond the constraints of traditional laboratory settings.

## Introduction

Ambulatory assessment utilizes diverse data sources to gain a deeper understanding of people’s thoughts, emotions, and actions in their natural environment [1]. The simultaneous acquisition of physiological, behavioral, and self-report data allows the study of individual behavior under natural conditions [2, 3]. In contrast to traditional and highly controlled laboratory-centric approaches, ambulatory assessment methods can be systematically integrated into real-life environments through which individuals naturally navigate. This involves the use of portable devices and wearable sensors to capture continuous data streams in real-time [1]. Such approaches attempt to maximize ecological validity as they enable to automatically assess environmental variables (e.g., temperatures, geolocations, time of the day; [4]) while also capturing the temporal dynamics and contextual intricacies of psychological and subjective phenomena, as well as human behaviors, thereby minimizing potential biases associated with retrospective self-reports [5, 6]. Furthermore, the constraint to laboratory settings poses challenges in transferring findings to real-world scenarios, potentially limiting the generalizability of observed psychophysiological responses [7]. By incorporating multimodal psychophysiological measures, these approaches can improve our understanding of the reciprocal interactions between psychological processes and physiological responses as they occur under natural conditions [8].

Activity of the autonomic nervous system (ANS) as a proxy of the internal arousal state, plays a pivotal role in numerous behaviors and emotions [9, 10], operating independently of conscious effort and beyond voluntary control [11]. The ANS comprises two distinct components, the sympathetic nervous system (SNS) and the parasympathetic nervous system (PNS), which function as physiological antagonists, synergistically, or independently [12]. Commonly used physiological indices include cardiovascular activity (e.g. heart rate, heat rate variability), respiratory, and electrodermal activity (EDA). EDA is a direct measure of the current conductive properties of the skin [13] and is closely linked to specifically SNS activity [13, 14]. Heart rate variability, which measures variability in the time interval between successive heart beats, is closely related to PNS activity or vagal tone [15, 16]. Heart rate and skin conductance level tend to increase with increasing arousal. Concurrently, heart rate variability is being increasingly recognized as an indicator of autonomic flexibility and the ability to regulate emotions [15]. A combination of these at least partly complementary measures appears promising when aiming to comprehensively characterize the internal physiological response associated with emotional processing [17, 18], or self-regulatory mechanisms linked to cognitive, affective, social, and health phenomena [19, 20].

Various laboratory studies found that measures of ANS signaling convey objective information related to stimulus processing and decision-making [21–23], that is not captured by subjective ratings [24, 25]. Physiological measures can therefore offer valuable insights to (clinical) psychology [26–28], and into how individuals respond to different contextual factors [25, 29, 30]. Natural environments, influencing emotional well-being and stress [31, 32], also utilize psychophysiological measures in clinical studies [33, 34], with evidence indicating positive effects on psychosocial well-being [35, 36], restoration, cognitive function [37], heart rate variability [38, 39], and brain activity [40] from walking in urban green spaces. Ambulatory psychophysiological assessment approaches also have potential utility for real-time monitoring and intervention in substance use disorders [34]. Reliability assessments of psychophysiological metrics have largely focused on data obtained in controlled laboratory settings. Here, heart rate and heart rate variability exhibit moderate to excellent reliabilities [22, 41–43]. Likewise, lab-based EDA measures show moderate reliability [13, 44–46]. In contrast, there is a dearth of studies investigating the reliability of ambulatory measures in naturalistic settings, particularly with regard to different physiological metrics. Although studies specifically focusing on the reliability of various physiological metrics in urban naturalistic contexts are lacking, some studies have reported the reliability estimates of physiological parameters during exposure to outdoor environments (e.g. nature viewing, outdoor walks, and outdoor exercise) [47]. Cardiovascular indices like heart rate and blood pressure have shown acceptable reliability during outdoor mobility [48–50]. However, there is a scarcity of evidence regarding the reliability of EDA measures in naturalistic conditions, as (to our best knowledge) no studies reporting reliability outcomes in outdoor environments have been found in the literature.

Bridging the gap between laboratory reliability and real-world settings, understanding how these metrics perform in naturalistic environments is critical to ensure the robustness of ambulatory psychophysiological measures in diverse settings [51]. The absence of substantiated evidence of reliable measurements in natural settings raises questions about their practical utility. Ensuring precision and consistency in psychophysiological measures is pivotal for robust conclusions regarding the link between exposure to natural settings and clinical research, necessitating rigorous testing and comprehensive validation of these instruments for generalizability in various ecological contexts [51, 52]. Ambulatory assessment approaches come with unique challenges that might impact reliability. Physiological signals are obtained across a wide variety of environments and bodily states [53], such that there is an inherent trade-off between reliability and validity [54]. Emphasizing reliability through standardized procedures and equipment settings can enhance consistency in measurements but may reduce validity by oversimplifying real-world phenomena. Conversely, improving ecological validity by sampling under a range of contextual settings undermines reliability [55–57]. Balancing the need for standardized replicable protocols to ensure reliability with the imperative to maintain the ecological validity of measures is crucial for mitigating this trade-off.

This study aims to navigate this balance, recognizing that decisions made to enhance the reliability of psychophysiological measures in real-world contexts inevitably influence their validity, and thus, necessitate a thoughtful and context-specific approach. Nonetheless, despite the limited extent of reliability testing for psychophysiological parameters in natural settings, a deliberate and transparent attempt as illustrated in this study, is clearly warranted. To the best of our knowledge, this study represents the preliminary investigation into the test-retest reliability of multiple psychophysiological data within the context of navigating a real-world urban environment. Here we addressed this issue by examining the reliability of physiological measures (electrodermal- and cardiovascular activity) in healthy individuals when navigating an urban route under naturalistic conditions. Psychophysiological activity and outside temperature were recorded using mobile sensors on two separate testing days, and movement and location were tracked via global positioning system (GPS). The data were obtained in the context of a larger study exploring metabolic influences, such that participants were tested once in a fasted and once in a sated state. This allowed us to examine whether satiety had a general impact on physiological measures. Overall, the test-retest reliability of a compound score across physiological measures (obtained via principal component analysis) exceed that of most individual measure and was acceptable. Results show that even under highly uncontrolled naturalistic conditions, ambulatory physiological measures exhibit acceptable reliability.

## Methods

### Sample

Twenty-six healthy participants (*N* = 15 female, *Range:* 18–34 years) took part in the study. The sample size was selected based on practical and financial considerations (e.g. we had a time window of around 4 months for data acquisition). Inclusion criteria included no history of cardiovascular, metabolic, gastrointestinal, psychiatric, or neurological disease, no medication or drug abuse. Participants were not on a diet and did not change their eating habits during the testing period. Vegetarians (*N* = 6) and vegans (*N* = 2) were included in the sample. Participants’ body mass index (BMI) was in the normal range (19.0 to 26.4 kg/m², *M* = 21.94, *SD* = 2.19), as was the percent body fat (*M*_female_ = 21.86, *SD* = 7.18; *M*_male_ = 14.72, *SD* = 4.57). All participants provided written informed consent and received a reimbursement of 10€ per hour for their participation. The study was approved by the ethics committee of the University of Cologne (code: JPHF0120).

### General Procedure

The data in this paper stem from a larger two-day study, assessing the effect metabolic state (sated vs. fasted condition) on various measures. Employing a counterbalanced within-subjects design, participants underwent two separate sessions on separate days, with a 6–8-day interval in between. On each testing day, participants walked the same urban route while undergoing an ambulatory psychophysiological assessment. The entire procedure was performed once in a sated state and once in a fasted state, in counterbalanced order. In the fasted state, participants abstained from food intake for approximately 10 hours before the testing session, while in the sated state, participants consumed breakfast prior to the session.

On the first testing day, participants also underwent a baseline screening in the laboratory to check exclusion criteria. Additionally, information regarding dietary preferences such as veganism or lactose intolerance was collected. A Bioelectrical Impedance Analysis to assess BMI was performed for each participant, followed by the completion of several questionnaires, including socio-demographic details and subjective assessment of tiredness and hunger levels. Participants were then provided with a detailed map of the urban walking route and were instructed to memorize it after a thorough explanation. The route included three intersections with traffic lights, a small path between two green areas and an avenue of trees leading to and from the main street (see Figure 1). Subsequently, mobile electrocardiograph (ECG) and EDA sensors were applied (see next section). A study smartphone equipped with GPS recording capabilities was provided to the participant. Communication with other people or listening to music was prohibited during the urban walking exposure. Participants were instructed to walk at a constant, previously trained speed (*M* = 4.1 km/h, *SD* = 0.27). The urban route was 2.1 km in length and took participants approximately 30 minutes to complete. Upon return, GPS recording was terminated, and electrodes were removed from the participant in the laboratory. Participants were asked about any noteworthy events they may have experienced during the urban walking route and the experimenter noted them accordingly. After route completion participants completed an additional laboratory task, to assesses subjective evaluation of food rewards.

**Figure 1.**
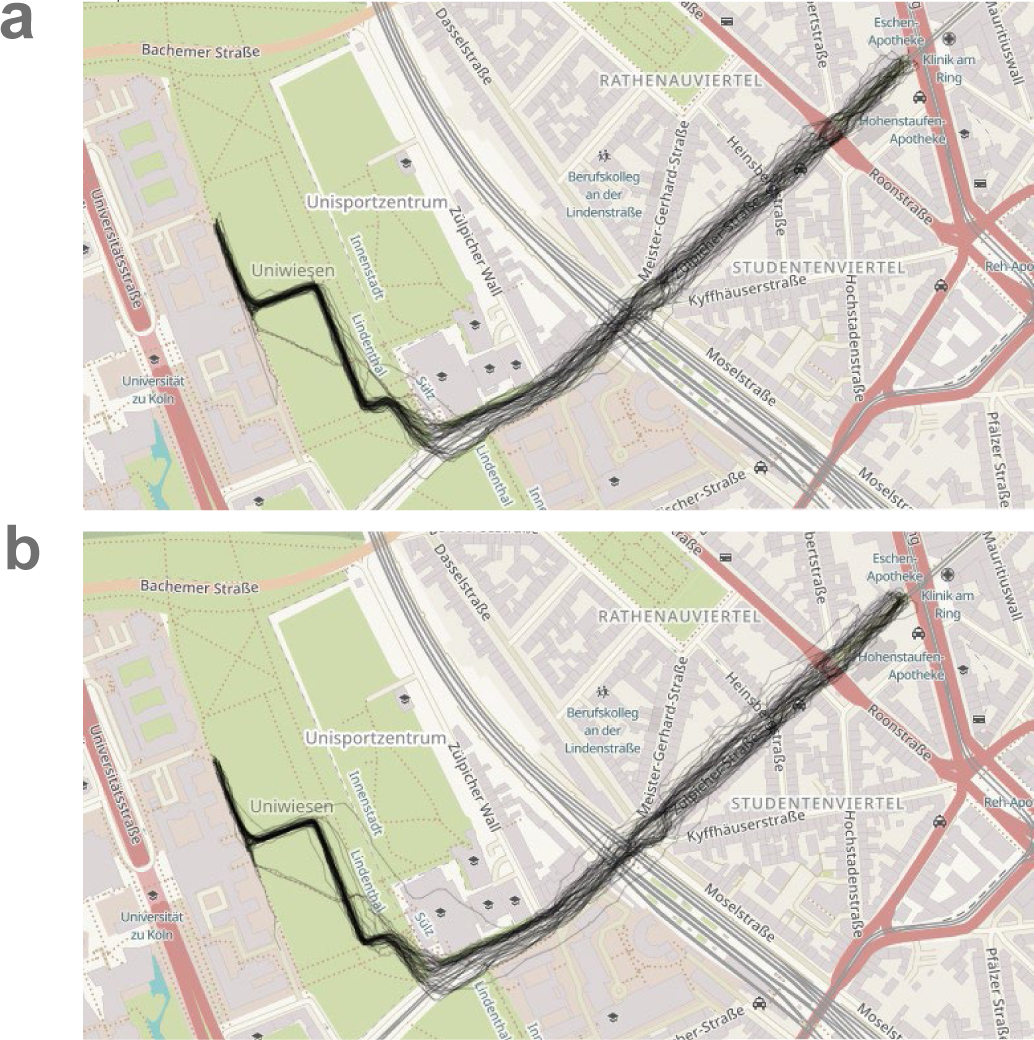
Urban walking route with single-subject trajectories. Urban walking route for day one (a) and day two (b). Individual-participant movement trajectories derived from GPS-based location tracking are depicted by black lines. Visualization created using leaflet-package [69].

### Ambulatory assessment

#### Physiological recording

To measure ANS activity, electrodermal and cardiovascular data were recorded using the mobile ECGmove4© and EDAmove4© devices from Movisens [58]. Both devices store data in terms of the system time of the associated computer they were coupled with, ensuring synchronization of time stamps across channels. The ECGmove4© was placed under the left chest of the participant using disposable adhesive pre-gelled Ag/AgCI electrodes and acquired raw data (sampling rate 2000 Hz) of a single channel ECG, from which secondary parameters like heart rate variability (ms) and heart rate (bpm) were calculated. The EDAmove4© collected skin conductance level (µS), skin conductance response (count), and skin conductance response-amplitude (µS) data to reflect EDA measures. The EDA device was fixed to a strap band on the non-dominant wrist. The EDA pre-gelled disposable adhesive Ag/ AgCI electrodes were attached to the thenar and hypothenar eminences of the non-dominant palm. Further, the sensor applied a DC voltage of 0.5 V to the skin to gain skin conductance with a 32 Hz sampling rate.

#### GPS-based location tracking

Participants were equipped with a study smartphone for continuous GPS-location tracking. Time and date on the GPS location tracking app (GPS Logger version 2.1.12) were synchronized with the mobile physiological sensors via the linked computer. The tracking of the instantaneous position was performed at an interval of one second to ensure high accuracy.

### Data analysis

#### Physiological data

Statistical analyses were carried out using R Studio © 2024a [59] and Matlab © 2019a (MathWorks) statistical software [60]. Physiological data were processed using the custom Movisense © “DataAnalyzer”-software and using custom Matlab © code. Generally signal quality was assessed via visual inspection, preprocessing, correction for artifacts from motion, and involved an alignment of recordings to the 30-minute urban walking route. Preprocessing of the raw EDA data involved a lowpass filter set at 0.1 μS to correct high frequency noise and related artifacts. Raw ECG data underwent a down-sampling process, transitioning from an initial acquisition frequency of 2000 Hz to 200 Hz. Down-sampling is a standard procedure in signal processing, allowing for more efficient analysis while retaining essential information. Due to missing values in heart rate variability indices, a custom Matlab © code was developed to identify R-peaks from the raw ECG data for heart rate variability analysis. To enhance the accuracy of peak detection, the ECG data was first z-standardized. Subsequently, a threshold of 0.8 Hz was established. This threshold serves as a criterion for identifying significant peaks in the ECG signal. The purpose of setting this threshold is to filter out noise and ensure that only prominent peaks, which are indicative of R-waves, are considered during the analysis. By employing this threshold, we aim to improve the reliability and precision of the heart rate variability analysis results. Subsequently, the variance between R-R intervals was computed as the Root Mean Square of Successive Differences (RMSSD), a standard time-domain heart rate variability metric [43, 61].

#### Data processing

For each participant and physiological parameter of interest (heart rate, skin conductance level, skin conductance response amplitude, and skin conductance response count) physiological measures were computed for successive sixty second time bins, covering the entire 30-minute route. Furthermore, physiological measures were segmented into five recording epochs, including baseline-urban green space (park), three segments of urban grey space (exposure to shops and restaurants), and post-urban green space (park). To achieve this, GPS data were co-registered with the mobile sensors time stamps, ensuring consistent temporal alignment. The route with predefined start and end coordinates (latitude: 50.9272495, longitude: 6.9343163) served as a reference for epoch delineation. Participants movement along this route was tracked, with the initiation of the urban grey space phase marked by proximity to these coordinates and its conclusions indicated by deviation from the specified route. Subsequently, the urban grey space was subdivided into three equal segments. Following segmentation, physiological measures were averaged per segment, and z-standardized within-participants, separately for each testing day. In addition, in analyzing the comprehensive physiological responses along the entire walking route, the physiological measures were averaged across all epochs for each participant and each testing day, covering the entirety of the walking route. We subdivided the physiological measures collected during the study into two distinct partitions. The first partition segmented data into five segments. The second partitioning used the average physiological data across the entire recording epoch for each participant and session, providing a comprehensive overview of physiological responses throughout the entire walking route. These partitions offer complementary perspectives on the temporal dynamics of physiological activity during the study.

#### Principal Component Analysis

To account for covariance between physiological measures, we applied Principal Component Analysis (PCA). PCA is a multivariate statistical technique for data-driven dimensionality reduction that identifies directions of shared variance in high-dimensional data. PCA identifies linear combinations of the original variables that project the data onto orthogonal axes. Based on two distinct partitioning’s derived from the physiological measurements (as described above), PCA was conducted separately for each partitioning. For the first partition, separate PCAs were computed for each segment of the data, per day. In contrast, for the partition covering the entire walking route, a single PCA was computed across all physiological measure, separately for each day. This approach captured variations in physiological responses over both testing days, facilitating exploration of underlying patterns.

#### Reliability assessment

Test-retest reliability of principal component scores and individual physiological measures were computed using a bootstrapping approach. For each measure, 10k bootstrap samples were created via resampling, and the Spearman (*r*_S_) correlation coefficient (correlation between day one and day two measures) was computed for each bootstrap sample. The median of the correlation coefficient distribution across samples was taken as a summary measure. The reliability assessments followed established criteria, recognizing that reliability estimates are continuous measures [62]. Rather than relying on arbitrary cutoffs, some researchers suggest labeling intervals of reliability [63]. Reliabilities were classified as follows: *r* ≤ 0.30: low; 0.31≤ *r* ≤ 0.50: moderate; 0.51≤ *r* ≤ 0.70: acceptable [64], 0.71≤ *r* ≤ 0.80: good [65], 0.81≤ *r* ≤ 0.90: very good [66], *r* > 0.90: excellent [67]. Note that such labels provide additional context for interpreting reliability estimates, and should not be viewed as strict thresholds [68].

### Data availability

Raw individual-subject data will be made available upon publication via the Open Science Framework.

## Results

GPS tracking confirmed that participants consistently adhered to the prescribed walking route (see Figure 1a and 1b for individual participant movement trajectories). Throughout the study, environmental temperatures ranged between 12° and 37 ° degrees Celsius (*M* = 19.8°).

As a first step, we visualized trajectories of individual mean physiological measurements across the route (averaged for bins of sixty seconds). On both testing days (see Figure 2), physiological measures consistently fell within plausible ranges, with considerable variability across participants.

**Figure 2.**
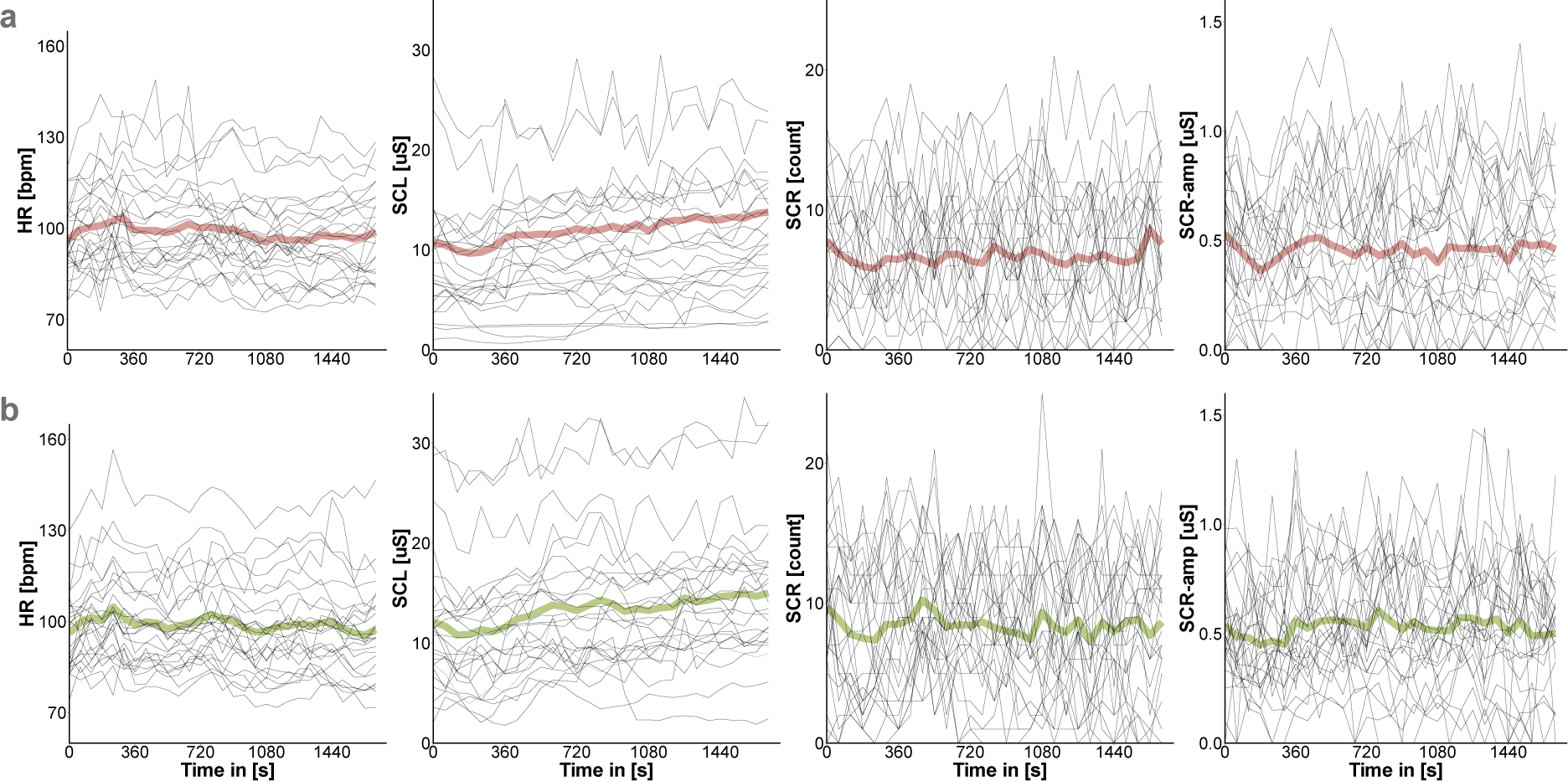
Physiological measures for single individual participants (thin lines, *N*=26) and group averages (thick lines) across time for day one (a) and day two (b). HR = Heart Rate, HRV = Heart Rate Variability, SCL= Skin Conductance Level, SCR-amp = Skin Conductance Response-Amplitude, SCR = Skin Conductance Response count.

Next, we separated data according to testing day (day one vs. day two, see Figure 3a) and condition (sated vs. fasted, see Figure 3b), to explore potential order and/or metabolic modulation effects.

**Figure 3.**
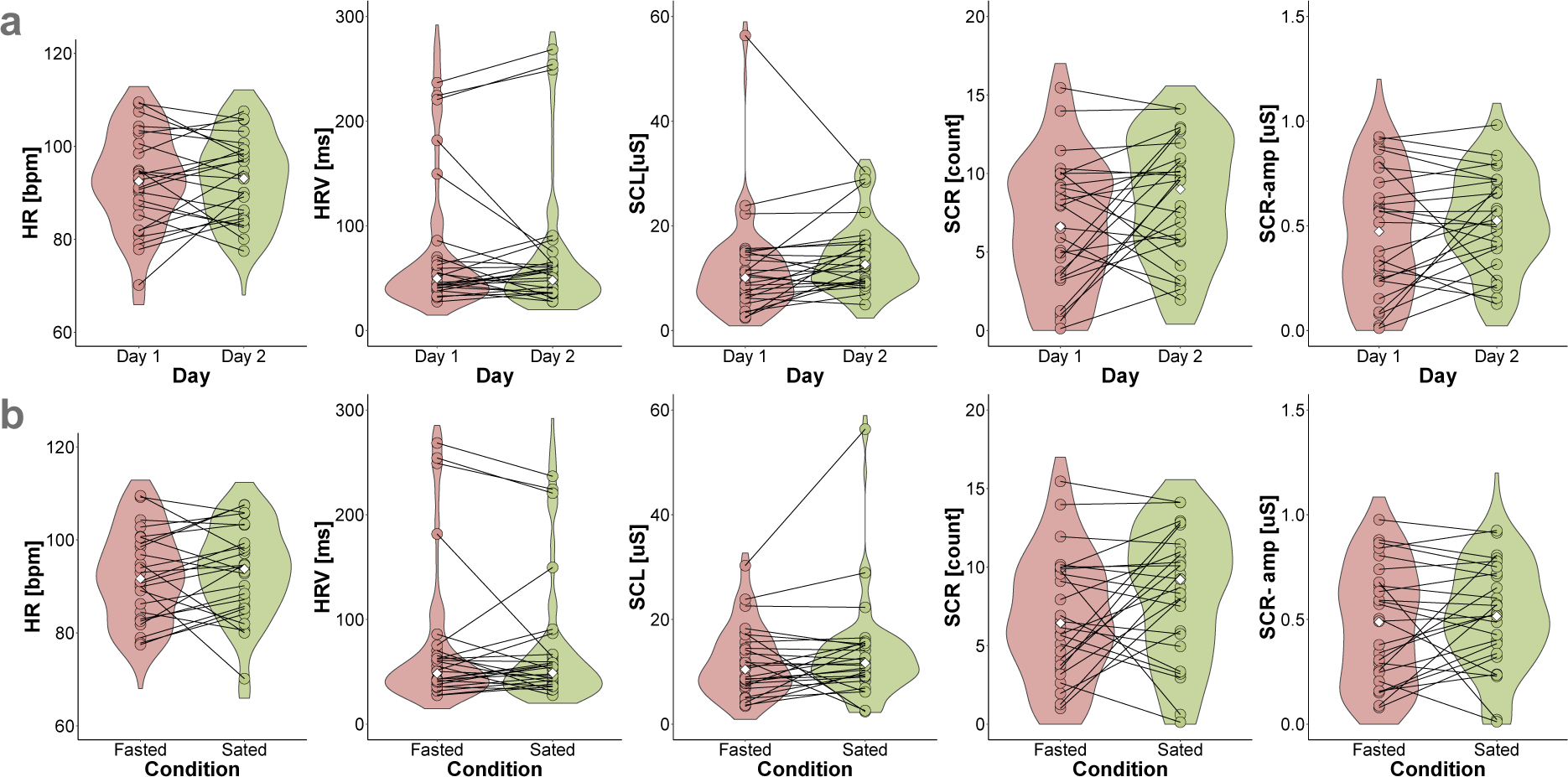
Distributions of mean physiological measures across days and conditions. Violin plots depicting mean physiological data from both testing days (panel a) and from both conditions (panel b); dots represent SS means of each day/condition; white diamond mark = median. HR = Heart Rate, HRV = Heart Rate Variability, SCL= Skin Conductance Level, SCR-amp = Skin Conductance Response-Amplitude, SCR = Skin Conductance Response count.

Differences in single physiological measures between testing days (day one vs. day two) and conditions (fasted vs. sated) were assessed via Wilcoxon signed rank tests adjusted for multiple comparisons using Bonferroni correction. None of the comparisons were significant (see Table 1). As expected, there was considerable covariance in physiological measures, and the overall pattern was highly consistent across day one (Table 2) and day two (Table 3) for Pearson bivariate correlations of single physiological measures and temperature. None of the physiological measures showed significant associations with environment temperature.

**Table 1.**
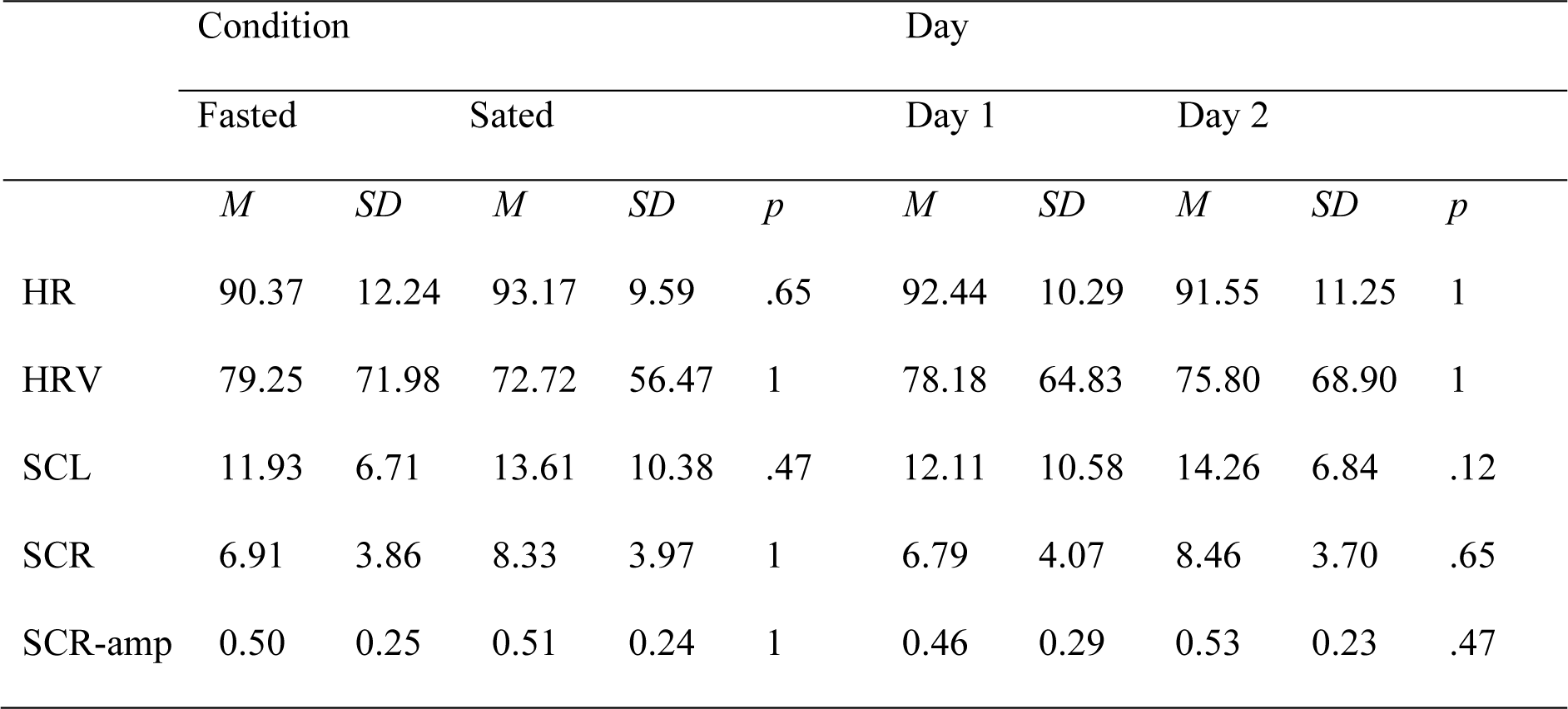
Wilcoxon signed-rank test of single physiological measures across conditions and testing days Bonferroni corrected. *N* = 26. HR = Heart Rate, HRV = Heart Rate Variability, SCL= Skin Conductance Level, SCR-amp = Skin Conductance Response-Amplitude, SCR = Skin Conductance Response count.

|  | Condition |  |  |  |  | Day |  |  |  |  |
| --- | --- | --- | --- | --- | --- | --- | --- | --- | --- | --- |
| | Fasted | | Sated | | $p$ | Day 1 | | Day 2 | | $p$ |
| | $M$ | $SD$ | $M$ | $SD$ | | $M$ | $SD$ | $M$ | $SD$ | |
| HR | 90.37 | 12.24 | 93.17 | 9.59 | .65 | 92.44 | 10.29 | 91.55 | 11.25 | 1 |
| HRV | 79.25 | 71.98 | 72.72 | 56.47 | 1 | 78.18 | 64.83 | 75.80 | 68.90 | 1 |
| SCL | 11.93 | 6.71 | 13.61 | 10.38 | .47 | 12.11 | 10.58 | 14.26 | 6.84 | .12 |
| SCR | 6.91 | 3.86 | 8.33 | 3.97 | 1 | 6.79 | 4.07 | 8.46 | 3.70 | .65 |
| SCR-amp | 0.50 | 0.25 | 0.51 | 0.24 | 1 | 0.46 | 0.29 | 0.53 | 0.23 | .47 |

**Table 2.** Pearson bivariate correlation matrix of single physiological measures and temperature on day one. Adjustment for multiple comparisons: Bonferroni. *N*=26, Correlation significant at \**p* > .05. HR = Heart Rate, HRV = Heart Rate Variability, SCL= Skin Conductance Level, SCR-amp = Skin Conductance Response-Amplitude, SCR = Skin Conductance Response count, Temp = Temperature.

| Variables | HR | HRV | SCL | SCR-amp | SCR | Temp |
| --- | --- | --- | --- | --- | --- | --- |
| HR | - | -.222 | .263 | .289 | .281 | -.213 |
| HRV |  | - | -.032* | .175 | .226 | .106 |
| SCL |  |  | - | .715* | .195 | -.084 |
| SCR-amp |  |  |  | - | .634 | -.099 |
| SCR |  |  |  |  | - | -.011 |
| Temp |  |  |  |  |  | - |

**Table 3.** Pearson bivariate correlation matrix of single physiological measures and temperature on day two. Adjustment for multiple comparisons: Bonferroni. *N*=26, Correlation significant at \**p* > .05. HR = Heart Rate, HRV = Heart Rate Variability, SCL= Skin Conductance Level, SCR-amp = Skin Conductance Response-Amplitude, SCR = Skin Conductance Response count, Temp = Temperature.

| Variables | HR | HRV | SCL | SCR-amp | SCR | Temp |
| --- | --- | --- | --- | --- | --- | --- |
| HR | - | -.508 | -.156 | .055 | -.044 | .091 |
| HRV |  | - | 0.007 | -.332 | -.255 | -.190 |
| SCL |  |  | - | .626* | .217 | .087 |
| SCR-amp |  |  |  | - | .358 | .287 |
| SCR |  |  |  |  | - | .280 |
| Temp |  |  |  |  |  | - |

### Principal Component Analysis

First, the data were split into five recording epochs per day (see methods section). For each epoch, a PCA was computed across physiological measures, separately for day one (Figure 4a) and day two (Figure 4b). Coefficients of the first principal component (PC#1) were mostly consistent across the recording epochs for both days (see Figure 4). For example, heart rate and EDA measures exhibited predominantly negative loadings on the PC#1.

**Figure 4.**
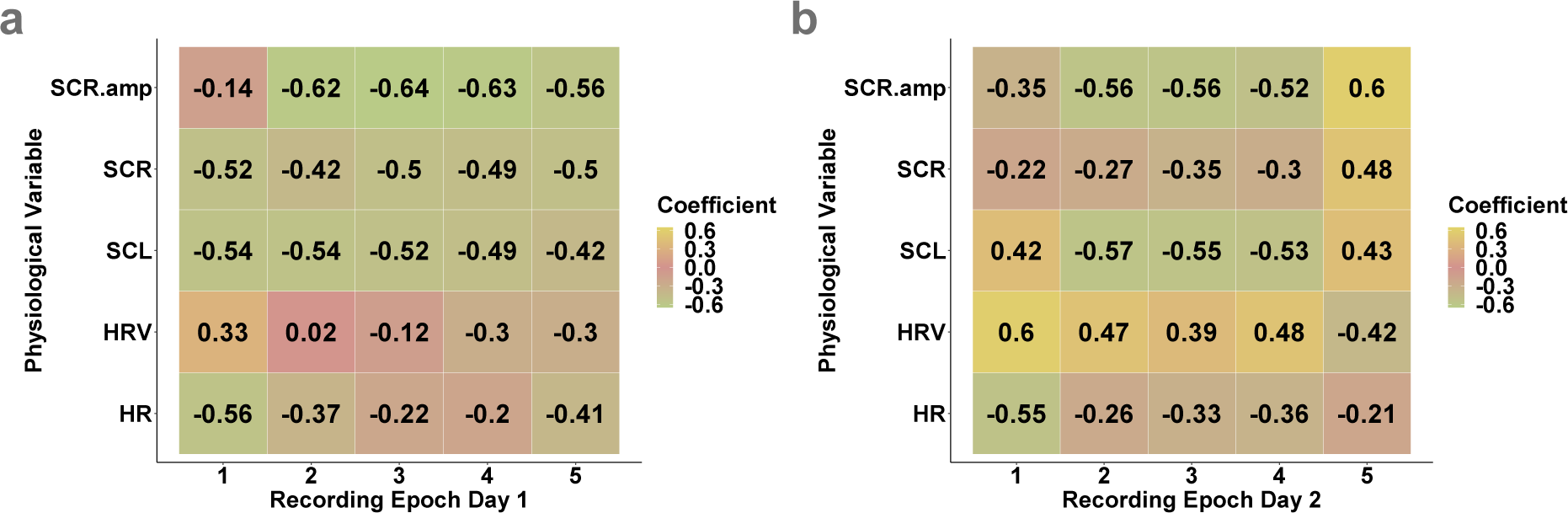
Heatmap of single physiological measures across recording epochs. Coefficients of PC#1 across recording epochs on day one (a) and day two (b). HR = Heart Rate, HRV = Heart Rate Variability, SCL= Skin Conductance Level, SCR-amp = Skin Conductance Response-Amplitude, SCR = Skin Conductance Response count.

Additionally, to comprehensively examine physiological responses throughout the walking route, the mean values of physiological measures were computed for each participant and testing day across all epochs. For each testing day a PCA across all physiological measures and participants was calculated. The PC#1 accounted for 39% to 45% of the variance across physiological measures on two distinct days (see Table 4). PC#1 coefficients were again highly consistent across recording days (see Table 5). This implies that the overall covariance pattern across physiological measures was stable across testing days.

**Table 4.**
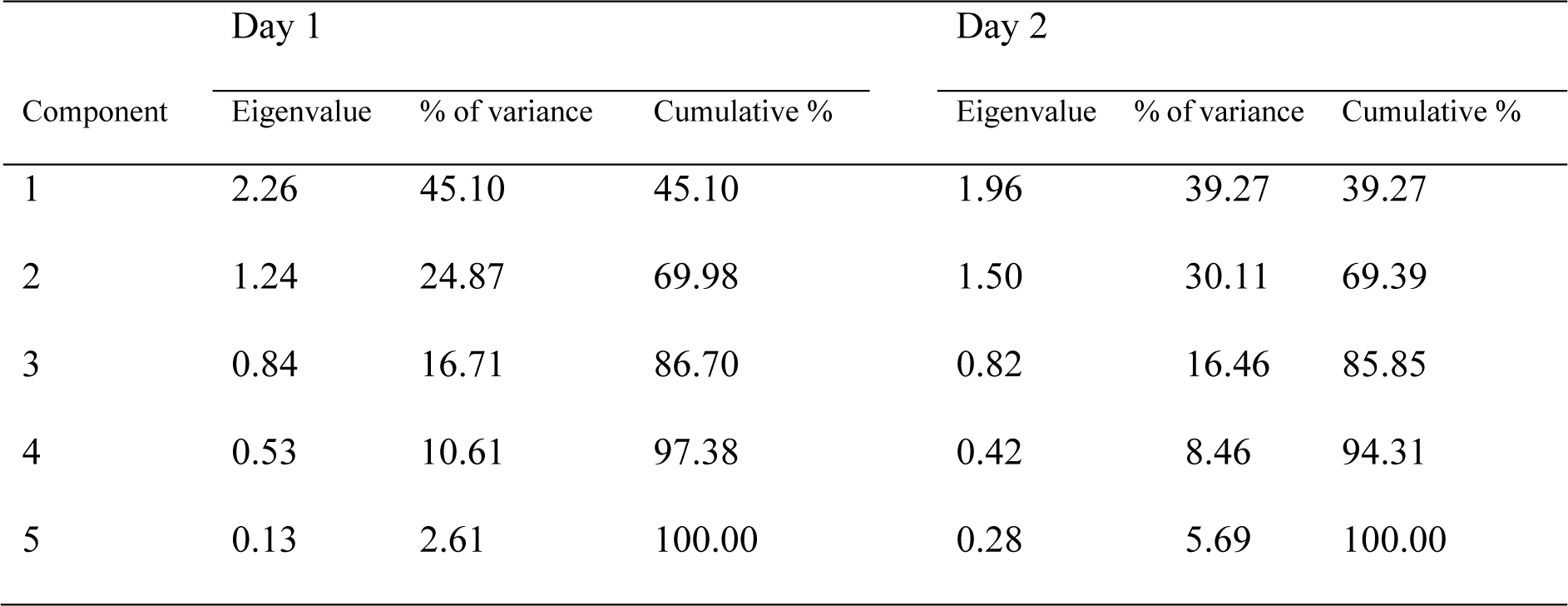
Contribution of the variance explained by all components, derived from the individual physiological measurements on different days across all epochs. HR = Heart Rate, HRV = Heart Rate Variability, SCL= Skin Conductance Level, SCR-amp = Skin Conductance Response-Amplitude, SCR = Skin Conductance Response count.

| Component | Day 1 |  |  | Day 2 |  |  |
| --- | --- | --- | --- | --- | --- | --- |
|  | Eigenvalue | % of variance | Cumulative % | Eigenvalue | % of variance | Cumulative % |
| 1 | 2.26 | 45.10 | 45.10 | 1.96 | 39.27 | 39.27 |
| 2 | 1.24 | 24.87 | 69.98 | 1.50 | 30.11 | 69.39 |
| 3 | 0.84 | 16.71 | 86.70 | 0.82 | 16.46 | 85.85 |
| 4 | 0.53 | 10.61 | 97.38 | 0.42 | 8.46 | 94.31 |
| 5 | 0.13 | 2.61 | 100.00 | 0.28 | 5.69 | 100.00 |

**Table 5.**
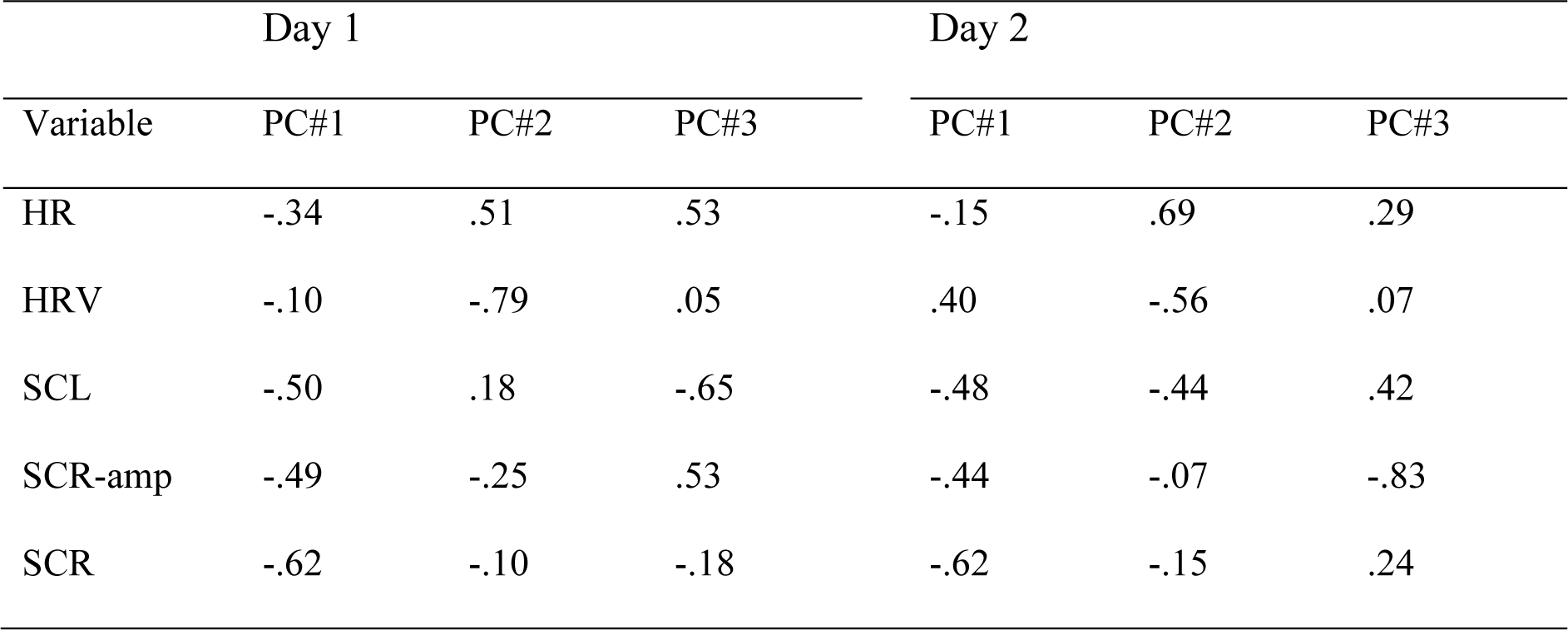
Coefficients of single physiological measures on different days across all epochs. Loadings of the first three principal components (PC#1, PC#2, PC#3) of the single physiological measurements at different days. HR = Heart Rate, HRV = Heart Rate Variability, SCL= Skin Conductance Level, SCR-amp = Skin Conductance Response-Amplitude, SCR = Skin Conductance Response count.

| Variable | Day 1 |  |  | Day 2 |  |  |
| --- | --- | --- | --- | --- | --- | --- |
|  | PC#1 | PC#2 | PC#3 | PC#1 | PC#2 | PC#3 |
| HR | -.34 | .51 | .53 | -.15 | .69 | .29 |
| HRV | -.10 | -.79 | .05 | .40 | -.56 | .07 |
| SCL | -.50 | .18 | -.65 | -.48 | -.44 | .42 |
| SCR-amp | -.49 | -.25 | .53 | -.44 | -.07 | -.83 |
| SCR | -.62 | -.10 | -.18 | -.62 | -.15 | .24 |

We next used a bootstrapping approach to estimate the test-retest reliability of each individual physiological measure as well as for the compound score of PC#1, across testing days (see Figure 5a-f). The median Spearman correlation coefficients across bootstrap samples indicate a low to acceptable reliability for individual physiological measures (HR: *r*_S_ = 0.53, HRV: *r*_S_ = 0.50, SCL: *r*_S_ = 0.53, SCR: *r*_S_ = 0.28, SCR-amplitude: *r*_S_ = 0.60). The reliability of PC#1 was acceptable (*r*_S_ = 0.60; see Figure 5f).

**Figure 5.**
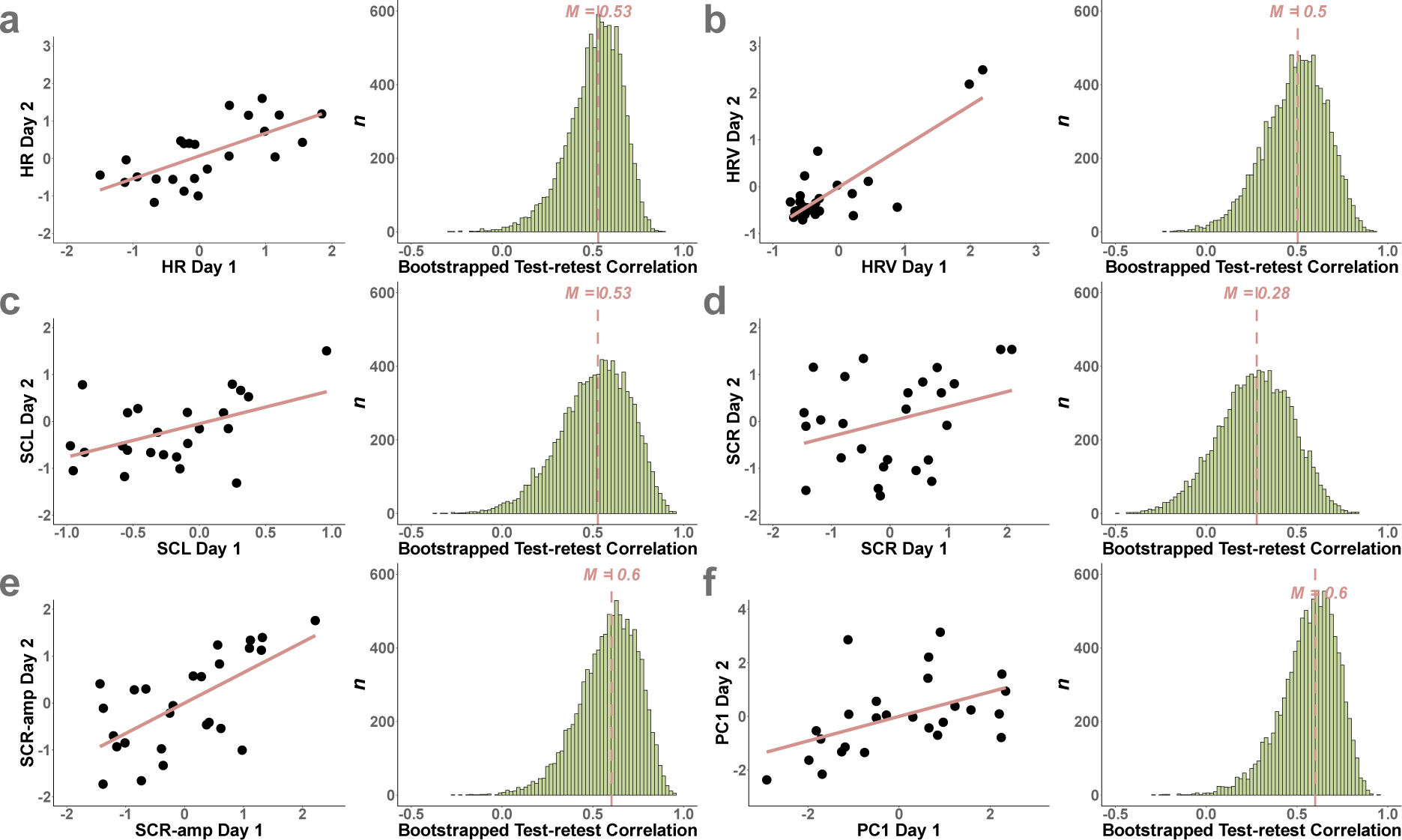
Bootstrapped test-retest analyses of single physiological data. Bootstrapped test-retest reliability of single physiological measures (a-e) over day 1 and day 2 (panel a) and bootstrapped test-retest correlation across 10k bootstrap samples (panel b); dashed red line: median correlation coefficient across samples. HR = Heart Rate, HRV = Heart Rate Variability, SCL= Skin Conductance Level, SCR-amp = Skin Conductance Response-Amplitude, SCR = Skin Conductance Response count.

## Discussion

In a counterbalanced repeated measures within-subjects design, we examined the test-retest reliability of ambulatory psychophysiological measures obtained under naturalistic ambulatory assessment conditions. Cardiovascular and electrodermal activity were continuously recorded using mobile sensors, in combination with GPS-based location tracking, while participants walked the same urban route on each testing day. The first principal component across physiological measures accounted for between 39% and 45% of variance across physiological measures. PC#1 scores from the aggregated measure exhibited acceptable test-retest reliability across testing days, suggesting that compound scores across physiological measures may provide reliable physiological indices under naturalistic conditions.

The loading pattern of PC#1 suggests that it captures the balance between parasympathetic and sympathetic tone, with highly similar patterns on both testing days. The negative association with most physiological measures suggests that higher scores on this component are indicative of reduced sympathetic tone. This interpretation is supported by the lower loadings observed for heart rate and EDA-based measures. Heart rate and heart rate variability tend to be inversely correlated [43, 70, 71]. The loading pattern observed for day two was consistent with this finding (i.e. PC#1 exhibited negative loadings for heart rate and positive loadings for heart rate variability). For testing day one, this pattern was less pronounced, and here heart rate variability showed a numerically small negative loading for PC#1. Nonetheless, the overall loading pattern for PC#1 was similar across testing days, and PC#1 score exhibited acceptable test-retest reliability across testing days.

In contrast, individual physiological measures demonstrated more variability in test-retest reliabilities, spanning a range from low to acceptable reliabilities. In particular the skin conductance response count measure exhibited low reliability. EDA measurements are especially susceptible to environmental fluctuations, movement artifacts [72], outside temperature and humidity [13]. Such effects may influence EDA measurements more than cardiovascular measures, in particular during ambulatory recordings. However, we did not observe associations of EDA measures with outside temperature, suggesting that this factor did not play a central role in the present study. The skin conductance level exhibited considerable variation among participants, ranging from minimal responses to significant reactions, consistent with the idea that this measure varies greatly inter-individually [14, 72]. Similarly, ECG-derived variables, such as heart rate variability, showed some outliers. This variability in some of the physiological measures highlights the utility of pooling information across them. Findings from the composite score (i.e. PC#1) indicate acceptable results in assessing participants’ physiological responses under naturalistic conditions, confirming the usefulness of combining information across multiple physiological measures in order to improve reliability.

Comparing the reliability estimates of physiological data obtained in controlled laboratory settings with those acquired under real-world, naturalistic conditions offers valuable insights into the robustness of physiological measures in diverse environments. Our findings indicate noteworthy similarities between the reliability estimates obtained in laboratory settings and those observed in naturalistic conditions. Specifically, cardiovascular indices such as heart rate and heart rate variability exhibit comparable moderate reliability estimates across both settings [41, 43]. For instance, Mathar et al. (2020) reported test-retest reliabilities (intraclass correlation coefficient) of 0.74 for heart rate and 0.78 for heart rate variability under laboratory conditions, showing a trend consistent with our findings of 0.53 and 0.50, respectively, under natural conditions. While our estimates exhibit a marginally lower reliability compared to those obtained in the laboratory, they nonetheless demonstrate a robust consistency. As such, the present results underscore a general robustness of cardiovascular indices across diverse environmental contexts and their efficacy as indicators of autonomic nervous system activity. Moreover, our investigation delved into the reliability of EDA measures, which are sensitive indices of sympathetic nervous system activation [13, 73]. Our results reveal that EDA measures maintain low to acceptable reliability levels during urban mobility. Notably, both skin conductance level and skin conductance response-amplitude demonstrate acceptable reliability estimates. Similar, laboratory studies show moderate reliability estimates of EDA measures [44–46]. While laboratory settings offer controlled conditions for data collection, our results suggest that physiological measures remain robust and reliable even in the face of environmental variability encountered in naturalistic settings. Moreover, our study highlights the potential advantages of employing a compound score derived from multiple physiological variables as a more comprehensive approach. By integrating data from various physiological measures, including cardiovascular and electrodermal indices, the compound score provides a summary representation of physiological response. This approach offers a more robust and nuanced understanding of participants’ reactions under naturalistic conditions, surpassing the limitations of single-variable analysis commonly employed in traditional lab-based research [5, 68].

We focused on reliability of psychophysiological measures during navigation in a naturalistic real-world environment. However, under such conditions, various factors might influence state-dependent changes in physiological measures. For instance, positive mood states are associated with an increased vagal influence on heart rate [75]. Baseline measurements and a GPS-based control of movement speed, as done in the present study, can help mitigate some of these effects. However, a 30-minute walk in a natural setting introduces the potential for unexpected stressors and/or mood fluctuations. Not all variations in mood and stressors can be fully accounted for, as participants may encounter unanticipated complexities during route exposure [51, 76]. While most participants did not report any incidents, some individuals reported specific experiences, such as encountering sirens or friends, or navigating dilemmas, all of which may potentially affect physiological responses. Such unsystematic effects likely lead to lower reliability estimates as compared to laboratory-based studies. However, put differently, despite the potential presence of uncontrolled factors, reliability was overall acceptable, in particular for the PCA-based compound score. By demonstrating the feasibility of obtaining reliable physiological data in real-world contexts, our study enhances the applicability and generalizability of physiological research, facilitating more ecologically valid assessments of human physiology. As this is a prerequisite for applications in (clinical) psychology, wearable health technologies, and digital health, the present results provide preliminary support for such future employment. Further research is, however, needed to delve into the specific factors driving potential patterns and their implications for human health and well-being.

### Limitations and future directions

As this study explored the reliability of physiological responses in a dynamic urban environment, several limitations emerged. First, we focused on the reliability of physiological measures across two testing days with a test-retest interval of around one week. Future research may delve deeper into the long-term stability of physiological measures. Second, the influence of external factors on physiological measures, such as environmental conditions (with the exception of temperature) were not controlled, such that the reliability estimates provided here constitute a lower bound – more tightly controlled experimental designs or recording conditions might yield higher estimates. Third, individual responses to specific stressors or events might reflect individual differences more than continuous physiological recordings such as carried out here. Future studies are required to explore this possibility. Fourth, we did not obtain subjective measures (e.g. stress or arousal ratings), such that the association between physiological and subjective measures remains unclear. Generally, future studies should replicate and test the reliability of psychophysiological data obtained in urban environments and/or under naturalistic conditions, and account for additional environmental effects and/or individual differences in response profiles. Future research might also further explore the associations of self-report measures and physiological effects.

## Conclusion

We examined the reliability of physiological measures obtained during naturalistic urban mobility conditions. Reliability of physiological measures is of central importance for studies that aim to shed light on how individuals navigate and respond to the complex interplay of environmental factors in natural settings. Our findings demonstrate an acceptable reliability of a PCA-based compound score across cardiovascular and electrodermal measures, which largely represented sympathetic tone during urban mobility. This confirms that physiological reactivity can reliably be estimated in a real-world urban setting, an important prerequisite for future applications of ambulatory psychophysiological assessments under real-world conditions. Contextual and environmental effects on physiological responses hold significant potential for ecologically valid future research applications in applied and clinical contexts in psychology and clinical neuroscience.

## Acknowledgements

J.P. acknowledges financial support for the Mapping Autonomic Neural Interaction and Control (MANIAC) Emerging Group by the University of Cologne Excellent Research Support Program via the German Research Foundation (DFG).

## Author Contributions

Conceptualization: Jan Peters & Kilian Knauth

Data curation: Pia Büning, Marie Schmeck, Dilber Korkmaz

Formal analysis: Dilber Korkmaz, Kilian Knauth

Funding acquisition: Jan Peters

Investigation: Pia Büning, Marie Schmeck, Kilian Knauth

Methodology: Dilber Korkmaz, Kilian Knauth, Jan Peters

Project administration: Jan Peters, Kilian Knauth

Resources: Jan Peters

Software: MATLAB 2019a, RStudio 2024

Supervision: Jan Peters

Validation: Kilian Knauth, Dilber Korkmaz

Visualization: Dilber Korkmaz

Writing – original draft: Dilber Korkmaz

Writing – review & editing: Jan Peters, Kilian Knauth, Angela Brands

